# The DOMINO web-server for active module identification analysis

**DOI:** 10.1101/2021.11.25.469692

**Authors:** Hagai Levi, Nima Rahmanian, Ran Elkon, Ron Shamir

**Author notes:** equal contribution (Ran Elkon, Ron Shamir).

## Abstract

Active module identification (AMI) is an essential step in many omics analyses. Such algorithms receive a gene network and a gene activity profile as input and report subnetworks that show significant over-representation of accrued activity signal (“active modules”). Such modules can point out key molecular processes in the analyzed biological conditions.

**Results:** We recently introduced a novel AMI algorithm called DOMINO, and demonstrated that it detects active modules that capture biological signals with markedly improved rate of empirical validation. Here, we provide an online server that executes DOMINO, making it more accessible and user-friendly. To help the interpretation of solutions, the server provides GO enrichment analysis, module visualizations, and accessible output formats for customized downstream analysis. It also enables running DOMINO with various gene identifiers of different organisms.

**Availability and implementation:** The server is available at http://domino.cs.tau.ac.il. Its codebase is available at https://github.com/Shamir-Lab.

## 1 Introduction

High-throughput omics data analysis frequently utilizes biological networks. In these networks each node represents a cellular subunit (e.g., a gene or its protein product) and each edge represents a relationship between two subunits (e.g., a physical interaction between two proteins). Integrated analysis of a biological network and a molecular profile measuring gene activity levels under a certain condition can greatly boost the functional interpretation of the data (Mitra *et al*, 2013). Activity levels can be calculated by measuring differential expression between two conditions or samples (Ideker *et al*, 2002; Chuang *et al*, 2007), by providing a set of genes associated with a disease as inferred from a GWAS (Nakka *et al*, 2016; Chang *et al*, 2015; Fernández-Tajes *et al*, 2019), or by estimating the mutation load of genes in cancer patients (Cerami *et al*, 2010). *Active Module Identification* (AMI) methods seek *“active modules”*, i.e., connected subnetworks that show a marked over-representation of high activity levels (Ideker *et al*, 2002; Leiserson *et al*, 2015). Such modules can reveal biological processes involved in the probed condition. A popular way to infer these biological processes is by conducting Gene Ontology (GO) enrichment analysis on each module (The Gene Ontology Consortium, 2019).

Recently, we evaluated six popular AMI algorithms and analyzed the GO terms that were enriched on their modules. We observed a high rate of non-specific calls of enriched GO terms in most algorithms, putting to question their capacity to illuminate processes that are specifically relevant to the probed conditions (Levi *et al*, 2021). Furthermore, we introduced DOMINO, a novel AMI algorithm with markedly higher rate of empirically validated calls (Levi *et al*, 2021). Of note, similar results were observed by a recent independent benchmark study, which also reported that DOMINO’s modules had substantially higher biological signals than modules found by other AMI algorithms (Lazareva *et al*, 2021).

The original DOMINO tool requires download and installation on the user’s machine. Here, in order to make DOMINO more accessible to researchers, we provide an online service requiring no installation. The server also enables GO-term enrichment analysis on each module, module visualization, standard output formats for downstream analysis, and options to run DOMINO with other organisms.

## 2 Materials and Methods

### 2.1 Input files: active gene sets and network file

The input for DOMINO is a set of active genes and a network. Note that DOMINO uses only binary gene scores (active/not active under the probed condition) and not real-valued scores. For the network, the user can upload a custom network file or choose a pre-loaded network. The available networks include DIP (Xenarios *et al*, 2002), HuRI (Luck *et al*, 2020) and STRING (Szklarczyk *et al*, 2017) with edge confidence score > 900. The preloaded networks use a cache mechanism (detailed in (Levi *et al*, 2021)) for faster runtime. In addition, the user can provide several active gene sets (e.g., for different diseases) in order to analyze and compare the results in a single execution.

### 2.2 resulting modules

After providing the input files and clicking the “execute” button, a request is sent to the server to run DOMINO and perform additional analyses. A typical execution takes up to two minutes. After execution, the resulting modules are visualized using Cytoscape.js(Franz *et al*, 2016). Genes contained in each module are annotated in the visualization. Alongside the module, the genes it comprises are shown. The user can navigate between different modules and solutions.

### 2.3 GO enrichment analysis

GO enrichment analysis is performed on each module and FDR corrected for multiple testing using the goaltools library (Klopfenstein *et al*, 2018). The background genes used for this analysis are those comprising the input network. A list of GO terms and their enrichment scores are reported in a table alongside the visualized module.

### 2.4 Downloading results

To enable further use of the solution, results can be downloaded by the user. Each module can be downloaded in two forms: (1) HTML (with the visualization and other results as they are shown in the DOMINO website), and (2) text files of the list of genes in the modules and a table summarizing the results of the GO enrichment analysis. This enables additional customized downstream analyses of modules and enriched GO terms.

### 2.5 Analyzing other organisms and gene identifiers

DOMINO uses by default ENSEMBL human identifiers. The website provides two options to run DOMINO with a list of non-human gene identifiers: (1) If the active gene list contains mouse ENSEMBL identifiers, and one of the pre-loaded networks is chosen, the genes in the active gene set will be converted to their corresponding human orthologs. In this case, GO enrichment analysis will be applied to the resulting modules. (2) If a custom network is supplied by the user, DOMINO matches the gene identifiers in the active gene list to the network. Note that in this case, the gene identifiers need not be taken from ENSEMBL, but can be of any species and GO enrichment analysis is not executed.

### 2.6 API calls for automated pipelines

To enable scripts to perform automatic calls to the server, we exposed a web-API for the execution of DOMINO. Details are provided on the landing page of the website.

## 3 Results

As a showcase, we uploaded an input set of 157 genes related to autism spectrum disorder (ASD) (taken from (Consortium, 2018)). We ran the tool with the preloaded STRING network. DOMINO detected eight modules in this run. Figure 1 shows the two largest modules. Reassuringly, they correspond to two distinct fundamental biological processes that are known to be severely abrogated in brains of ASD patients (Satterstrom. FK. *et al*, 2020): (1) chromatin remodeling and regulation of transcription (LaSalle, 2013; Cunniff *et al*, 2020) (Figure 1A,C) and (2) defects in neuronal trans-synaptic signaling (Guang *et al*, 2018)(Figure 1B,D).

**Figure 1:**
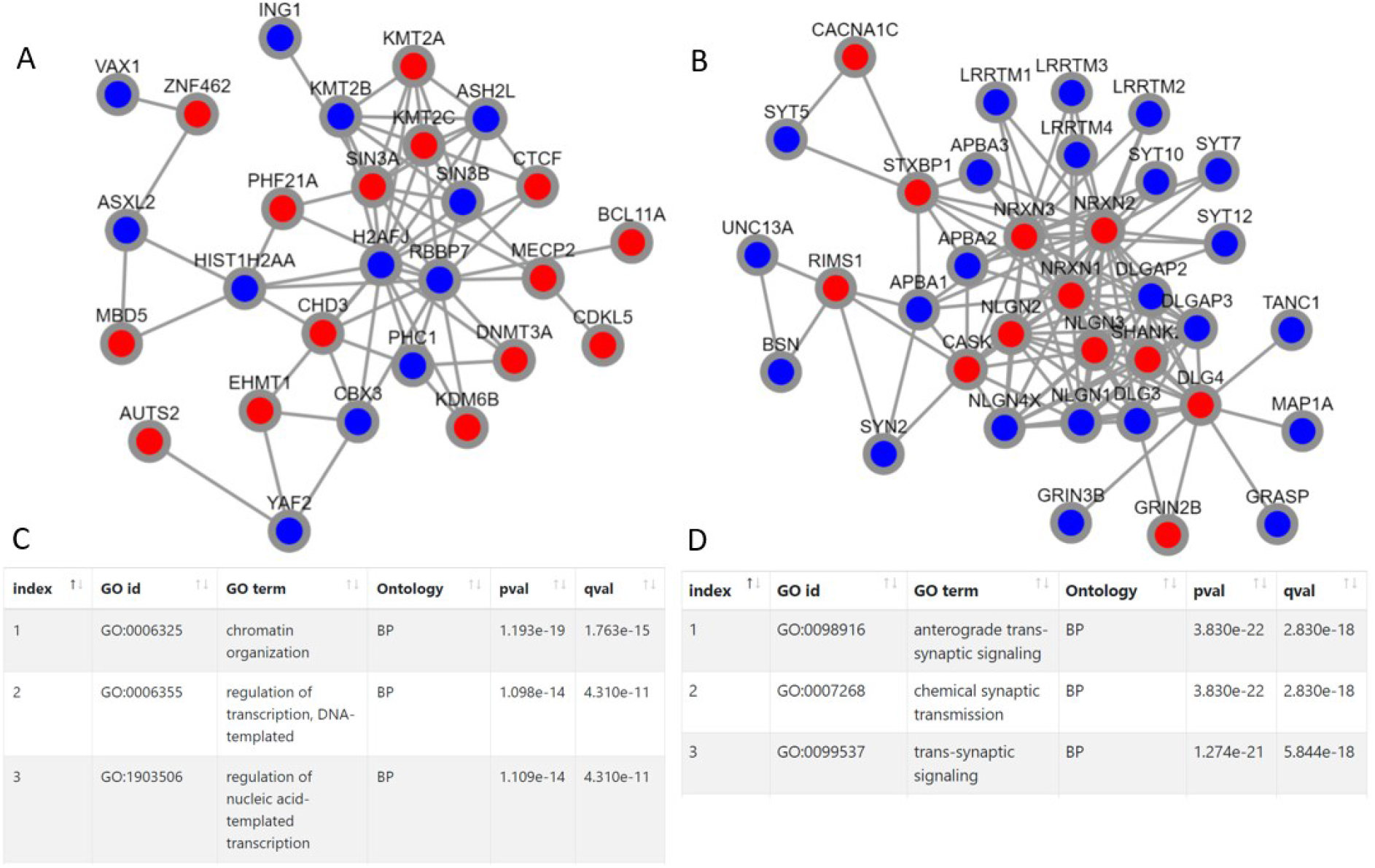
Two modules reported by DOMINO website on a set of 157 ASD related genes using the preloaded STRING network. The website runs the DOMINO algorithms and provides visualizations of the resulting modules (A, B) along with the most enriched GO terms found on each (C, D). The red nodes indicate the module’s genes that are included in the input set of active genes (here the set of ASD genes). Ontology: molecular function (MF), biological process (BP) or cellular component (CC); pval: nominal p-value; qval: FDR corrected p-value.

## Acknowledgements

Study supported in part by German-Israeli Project DFG RE 4193/1-1 (to RS and RE), by the Israel Science Foundation grant No. 1339/18 (to RS), by the Israel Science Foundation grant No. 2118/19 (to RE), by Len Blavatnik and the Blavatnik Family foundation (to RS) and the Koret-UC Berkeley-Tel Aviv University Initiative in Computational Biology and Bioinformatics (to RE and RS). HL was supported in part by a fellowship from the Edmond J. Safra Center for Bioinformatics at Tel-Aviv University. RE is a Faculty Fellow of the Edmond J. Safra Center for Bioinformatics at Tel Aviv University.

